# Programmable Repair of Disease-Causing UGA Stop Codons in Mammalian Brain

**DOI:** 10.64898/2026.05.13.724978

**Authors:** Ahmad Al Saneh, Lionel Gissot, Christopher A. Ahern

## Abstract

Protein truncating variants caused by UGA stop codons are the most prevalent class of rare variant mutations in neurodevelopmental diseases. Suppressor transfer RNA (sup-tRNA) have therapeutic potential for premature termination codon (PTC) repair, but have thus far underperformed by traditional AAV delivery platforms and progress has been hampered by the lack of methods to non-invasively assess in vivo activity in mammalian brain. To fill this material gap, we utilize transcranial in vivo bioluminescence imaging data from a luciferase-UGA mouse model to enable payload optimization. These data demonstrate that U6 promotor and AAV2/9 capsids have the lowest in vivo activity, whereas self-complementary AAV2/9 with the tRNA in a minimal 100bp genomic context provide broad and efficacious PTC rescue. Further, payload tRNA multiplexing and use of tRNA introns enable efficacy of low viral titers and sustained rescue. tRNA sequencing of scAAV delivered Arg^UGA^ sup-tRNA in brain demonstrate no effects on endogenous tRNA levels, their acylation or processing, and these features are also maintained in scAAV delivered Arg^UGA^ sup-tRNA. Collectively, this work defines a scalable strategy for precision UGA stop codon suppression, supporting development of durable genetic rescue therapies for neurodevelopmental disorders in the mammalian brain.

**GRAPHICAL ABSTRACT:** 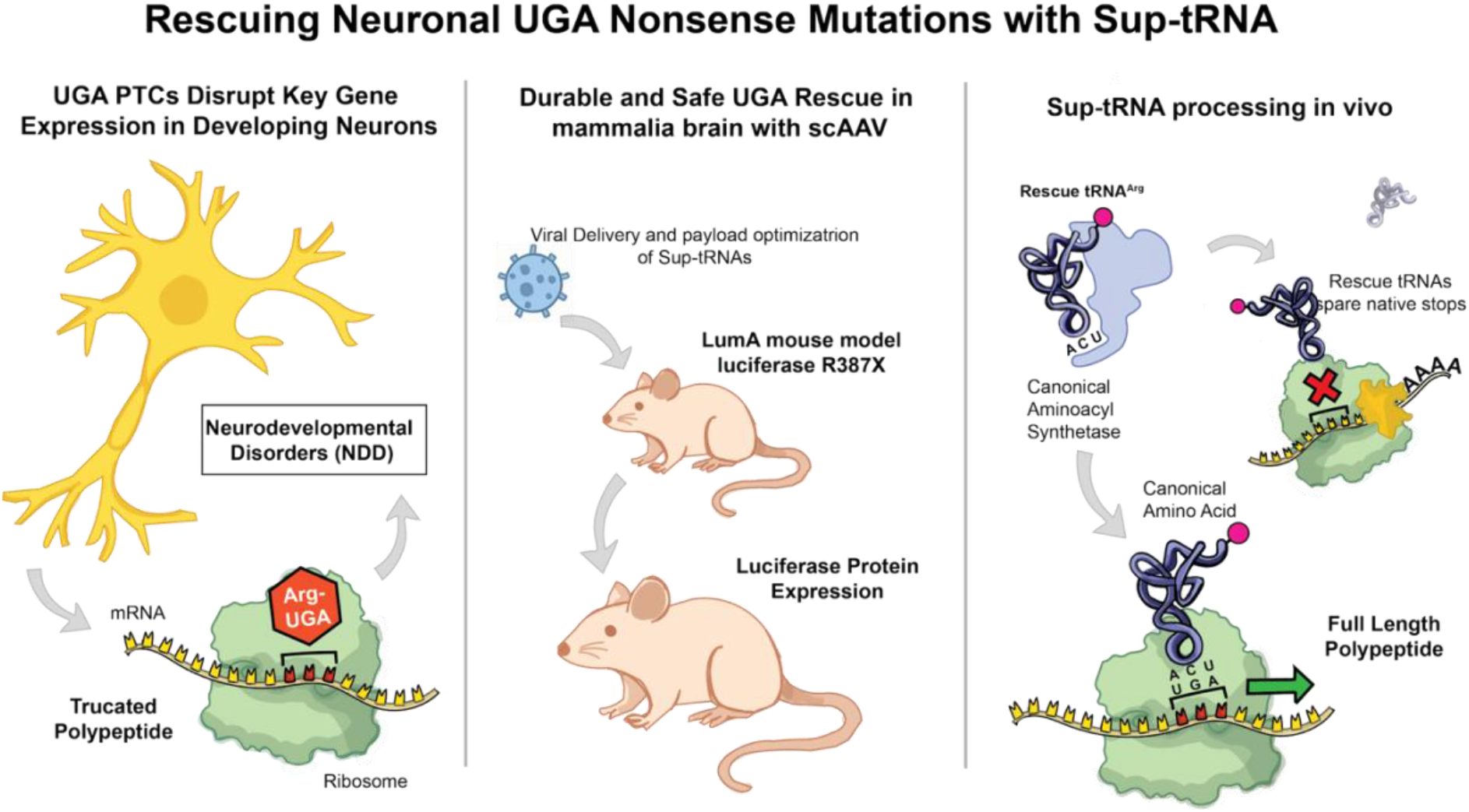

## INTRODUCTION

*De novo* protein truncations are common in neurodevelopmental disorders (1) and cause a range of genetic diseases, including intellectual disability (2), developmental delay (3,4), autism (5,6), epilepsy (7,8), as well as sleep and autonomic dysfunction (5,9). One type of truncating variant is caused by the creation of a premature termination codon (PTC), accounting for 10-15% of all inherited diseases (10). Such PTCs are nearly absent in population studies, indicating that they are closely associated with disease (1). While each PTC mutation is rare within any given disorder, a PTC can occur in principle throughout the entire coding sequence of a gene, thus posing a major therapeutic challenge. For this reason, efficient therapeutic strategies capable of repairing a given PTC in a sequence-agnostic manner are needed. Small-molecule read-through therapies have performed poorly in clinical trials (11), possibly due to their propensity to incorporate near-cognate amino acids at the PTC (12,13) and high off-target read-through of native stop codons (12–14). Additionally, while gene editing can correct a specific PTC, individual validation of guide-RNAs is necessary – a resource-intense effort that is currently not feasible for a multitude of rare (“N-of-1”) PTC variants. There is therefore a critical need for a broadly applicable treatment strategy that can correct PTCs, independent of the sequence context, with minimal toxicity in the brain. Notably, while ten amino acids (and 19 codons) are one nucleotide change away from becoming a PTC, the Arg CGA codon being mutated to the TGA stop codon represents the lion’s share of all PTCs (**Fig 1A**). This is due to a biochemical vulnerability in the CGA codon, where spontaneous deamination of cytosine to thymidine makes the CGA-to-TGA alternation the most common *de novo* PTC (15). Therefore, a single approach with the ability to rescue individual Arg-TGA PTC codons has broad clinical potential.

**Figure 1.**
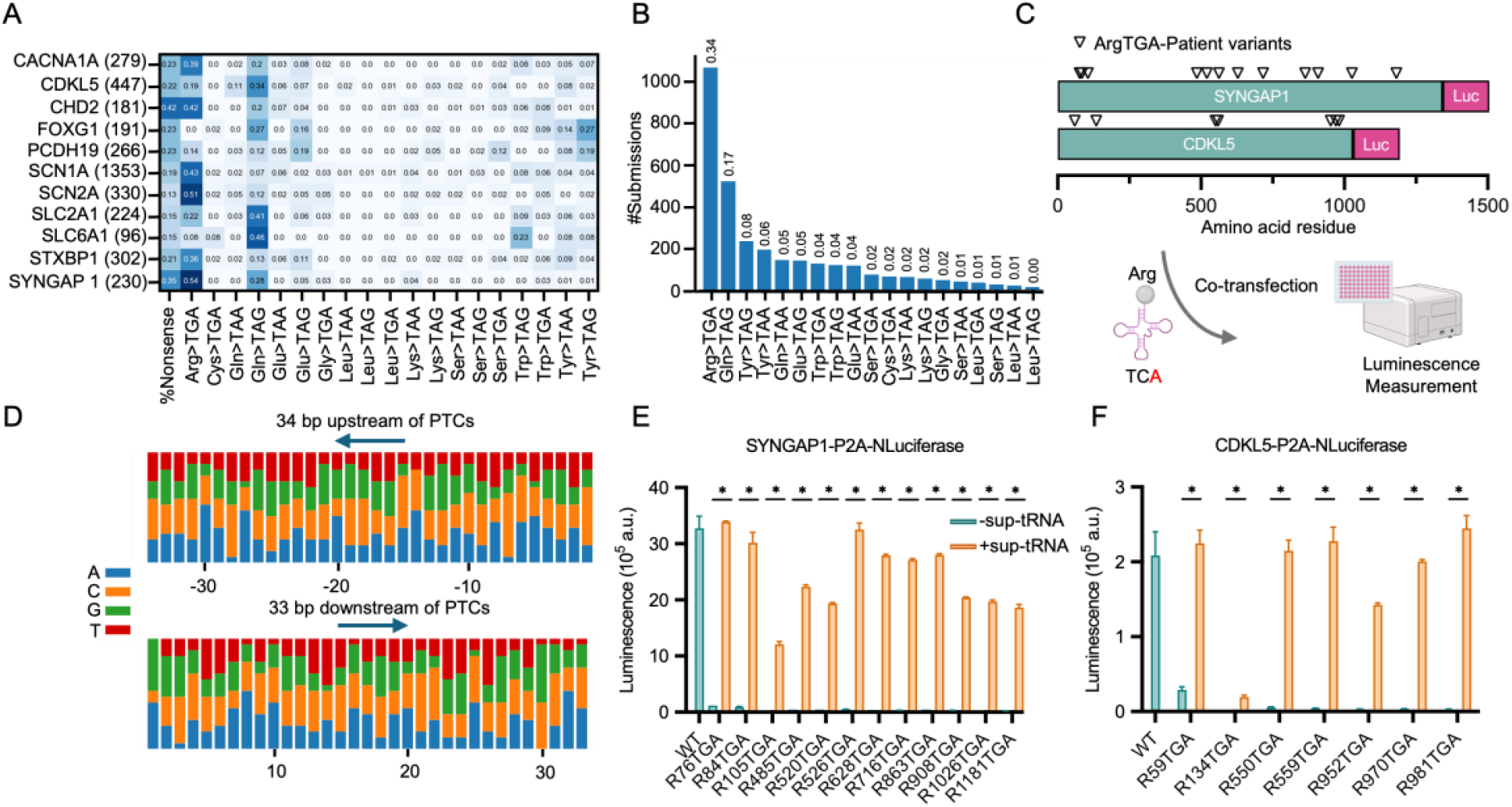
UGA variants of neurological disease and positionally agnostic rescue of UGA stop codons in SYNGAP1 and CDKL5. **(A)** Premature termination codon (PTC) landscape for a variety of neurological and neurodevelopmental disorders. Prevalence varies from %13 (*SCN2A*) to 42% (*CHD2*), with a majority caused by Arg CGA-to-TGA. **(B)** ClinVar submissions by stop codon type further demonstrates Arg^UGA^ as the dominant PTC. **(C)** experimental layout to measure agnostic Arg^UGA^ PTC rescue in *SYNGAP1* and *CDKL5*. **(D)** A 67-bp window centered on the mutated CGA codon (34 bp upstream and 33 bp downstream) across 19 variants from two genes: *SYNGAP1* (R76, R84, R105, R485, R520, R526, R628, R716, R863, R908, R1026 and R1181) and *CDKL5* (R59, R134, R550, R559, R952, R970 and R981). **(E-F)** positional agnostic rescue in *SYNGAP1* and *CDLK5*. Only one site, R134TGA in CDKL5, demonstrated poor rescue.

### Suppressor tRNAs have the potential to repair PTCs in a positionally agnostic manner

Suppressor tRNA (sup-tRNA) therapeutics have unique promise to promote protein restoration without introducing germline changes (13,16,17). We previously screened over 500 lab-engineered human sup-tRNAs to identify new variants that repair PTCs and encode the correct amino acid (18,19). These engineered tRNAs have been used for the rescue of PTCs in human hERG potassium channels (19), the cystic fibrosis transmembrane conductance regulator protein (20), CDKL5 (21), and *in vivo* for Hurler syndrome caused by PTC variants (22,23). This strategy is disease-modifying because it restores full-length protein from PTC-containing mRNAs that would otherwise undergo premature translational termination. Each sup-tRNA has a substitution in the anticodon triplet which allows reading of a stop codon, while still being able to deliver the correct amino acid (18). During translation, a sup-tRNA is accommodated by the ribosome at the PTC while delivering the correct amino acid to allow the ribosome to continue protein synthesis at that site and to proceed forward to codons past the PTC, generating a correctly translated and full-length protein. We and others have shown that this read-through of PTC does not obviously trigger read-through of native stop codons (18,24) or ER-stress (19,25), although more data are needed to parse the mechanistic basis for this phenomena. Recent efforts with virally delivered anticodon-edited arginine tRNA have focused on UGA stop codons that cause Hurler syndrome (Mucopolysaccharidosis type I, MPS I), however rescue in these cases remains subtherapeutic (<8% of WT protein) potentially reflecting limitations in AAV titers and/or payload designs. Sefl-complementary AAV (scAAV) vectors have shown some promise with recent rescue of retinal PTC variants (25), but no such example has been broadly described for neurological disorders. Lastly, there has been widespread concern regarding any tRNA-based therapy that could unintentionally and generally interfere with native tRNA processing and acylation, leading to potentially broad off-target consequences.

We have previously identified the anti-codon modified Arg-TCT-1-1 tRNA as one of the strongest amongst the 28 human arginine tRNAs (18,26). Specifically, the single anticodon mutation TCT-to-TCA reveals potent UGA suppression activity. We therefore first used sup-tRNA-Arg-TCA-1-1(hereafter, Arg^UGA^ sup-tRNA) to demonstrate positionally agnostic rescue of patient UGA variants in two genes, *SYNGAP1* and *CDKL5* while analyzing the sequence context of each PTC. We then adopt a transgenic mouse model carrying a *Rosa26* firefly luciferase gene harboring an Arg-to-UGA PTC (p. R387X) to optimize AAV and scAAV vector payloads containing the Arg^UGA^ sup-tRNA. This powerful approach allows non-invasive biodistribution and longitudinal analysis by transcranial in vivo bioluminescence imaging (IVIS). These data show broad sup-tRNA tolerance and activity, highlighting the requirement of placing the tRNA within a native and short genomic sequence (100bp) as well as the benefit of tRNA multiplexing. Strikingly, inclusion of tRNA introns increased scAAV titers by over 15-fold. Lastly, high-resolution tRNA sequencing performed on virally transduced brain samples reveals no consequence on native tRNA processing or expression, and shows that the anti-codon edited tRNA can achieve macroscopic rescue with expression levels at roughly ~2% of the native parental Arg-TCT-1-1 tRNA. This combined proteomic and in vivo vector-optimization strategy overcomes key barriers to clinical translation of suppressor tRNA therapeutics for monogenic brain disorders.

## MATERIAL AND METHODS

### Cloning of Arg^UGA^ sup-tRNA constructs for virus production

Four gBlocks gene fragments U6_tRNA_intron_Arg_TGA_TCT_1-1, U6_tRNA_Arg_TGA_TCT 1-1, Geno_noI_tRNA-Arg_TGA_TCT_1-1 and Geno_I_tRNA-Arg_TGA_TCT_1-1 (Supplementary Table 1) were designed with overlapping sequences (25bp), synthesized by IDT and cloned into the G0463 Self-Complementary Shuttle Plasmid from the University of Iowa Viral Vector Core (Ordering Adeno-Associated Virus Shuttle Plasmids | Viral Vector Core Facility - Carver College of Medicine | The University of Iowa) after linearization using the BamHI restriction enzyme. The gBlocks and the linearized plasmid were purified using Monarch® Spin DNA Gel Extraction Kit (NEB# T1120). The assembly reactions were performed using the NEBuilder® HiFi DNA Assembly Master Mix (NEB #E2621) according to the manufacturer’s protocol. The assembled plasmids were then transformed into stable3 chemically competent E. coli cells and plated on LB + Carbenicillin selective media. Positive clones were confirmed by colony PCR using LG153/154 primers (Supplementary Table 1) and by full plasmid sequencing at Plasmidsaurus.

The pCMV::mOrange2-Y156tga insert was PCR amplified using LG160/LG262 primers (Supplementary Table 1) and cloned into the G0463_pscAAV shuttle plasmid at the BamHI restriction site using NEBuilder® HiFi DNA Assembly, resulting in the pscAAV_pCMV::mOrange2-Y156tga-BgHpA plasmid. The four versions of tRNA-Arg-TCA-1-1 were PCR amplified using primers LG345/LG346 and LG206/207 and cloned into the pCMV::mOrange2-Y156tga-BgHpA plasmid at the SphI restriction site using NEBuilder® HiFi DNA Assembly.

### Cloning of SYNGAP1/CDKL5 Wild-Type and Arg-to-TGA Mutants

The full-length CDKL5 sequence (RefSeq: NM_003159, Clone ID: HsCD00022404) was obtained from the DNAsu repository at The Biodesign Institute, Arizona State University. The full-length SYNGAP1 sequence was kindly provided by Dr. Gavin Rumbaugh (University of Florida).

Wild-type coding sequences (CDS) of CDKL5 and SYNGAP1 were PCR-amplified using primers LG967/969 or LG935/937, respectively, and cloned into the pUC19_mcs_P2a_NanoLUC vector (linearized with KpnI) using NEBuilder® HiFi DNA Assembly, resulting in the constructs pUC19_CDKL5_P2a_NanoLUC and pUC19_SYNGAP1_P2a_NanoLUC.

To generate the Arg-to-TGA mutants, the mutant sequences were PCR-amplified using primer sets LG964/LG966/LG974–LG987 (for CDKL5) or LG964/LG966/LG1008–LG1038 (for SYNGAP1). These were cloned into pcDNA3.1 (linearized with BamHI and EcoRI) using NEBuilder® HiFi DNA Assembly, yielding pcDNA3.1::CDKL5_P2a_NanoLUC and pcDNA3.1::SYNGAP1_P2a_NanoLUC.

### PTC rescue in *SYNGAP1* and *CDKL5*

HEK293T cells were seeded in 6-well plates and cultured to approximately 80% confluency. Cells were then transiently transfected with 0.5 ~g of either pcDNA3.1::CDKL5_WT/TGAmut_P2a_NanoLUC or pcDNA3.1::SYNGAP1_WT/TGAmut_P2a_NanoLUC, along with 0.5 ~g of an ACE-tRNA-expressing plasmid, using PolyJet transfection reagent (SignaGen Laboratories). Twenty-four hours post-transfection, cells were lysed, and NanoLuciferase activity was measured using the Nano-Glo® Luciferase Assay System (Promega) on a Synergy Neo2 plate reader (BioTek).

### In vitro NanoLuc-TGA rescue

HT1080 cells were seeded in 96-well plates and transduced with the indicated sup-tRNA AAV vectors. At ~80% confluency, cells were transfected 6 h post-transduction with a NanoLuciferase reporter containing an in-frame TGA premature termination codon (NanoLuc-TGA) between Gly159 and Val160 using PolyJet transfection reagent, according to the manufacturer’s instructions. Luminescence was measured at 24 h and 48 h post-transduction using the Nano-Glo® Luciferase Assay System (Promega) and read on a Synergy Neo2 microplate reader (BioTek). Signal was quantified as relative luminescence units (RLU) for each well and analyzed as specified in the Data analysis/Statistics section.

### Animals

Heterozygous luciferase reporter mouse models (LumA) containing the R387X mutation (c.A1159T) in the luciferase gene located in the *Rosa26* locus of the mouse genome were generated by breeding homozygous reporter mice with wild-type (WT) mice. This breeding strategy ensured that all offspring were heterozygous for the luminescent transgene. Genotyping was performed at postnatal day 21 (P21) to confirm heterozygosity. All animal procedures were approved by the Institutional Animal Care and Use Committee (IACUC) and were conducted in accordance with institutional and NIH guidelines.

### Intracerebroventricular (ICV) Injections

Intracerebroventricular (ICV) injections were performed on postnatal day 0 to 1 (P0–P1) mouse pups. Neonates were anesthetized by hypothermia (placement on ice for ~3 minutes), in accordance with the institutional animal care and use guidelines (IACUC). Pups were secured in a Kopf neonatal stereotaxic frame and the head was stabilized for injection. Using a 32-gauge needle attached to a Hamilton syringe (PN: 7635-01), the specified dose of Arg^UGA^ sup-tRNA vector was delivered into the lateral ventricle using the following coordinates: (X, Y, Z) = (0.8, 1.5, 1.5) mm.

### In Vivo Bioluminescence Imaging

To assess in vivo brain luminescence, postnatal day 30 (P30) mice were imaged using the IVIS AMI imaging system (Advanced Molecular Imager, Spectral Instruments Imaging). D-Luciferin (GoldBio, Catalog Number: eLUCK-1G, CAS Number: 115144-35-9) was administered intraperitoneally at a dose of 150 mg/kg body weight. Following injection, mice were returned to their chamber and maintained under isoflurane anesthesia until imaging. A 20-minute interval was allowed to ensure sufficient penetration of luciferin across the blood-brain barrier. Imaging was performed with a 60-second exposure time and a field of view (FOV) set to 20 cm. Bioluminescent signal was quantified using AURA In Vivo Imaging Software (Spectral Instruments Imaging) by measuring photon flux within the region of interest (ROI) encompassing the brain.

### Ex Vivo Bioluminescence Imaging

For ex vivo analysis, mice were euthanized under isoflurane anesthesia, and brains were rapidly harvested and placed in small dishes. A total of 200 ~L of D-Luciferin solution was evenly pipetted over both hemispheres of each brain to ensure full substrate coverage.

Brains were imaged using the IVIS AMI system with ex vivo acquisition settings (60-second exposure). Luminescence was quantified by ROI-based photon flux measurements using AURA In Vivo Imaging Software.

### Sequences and Cloning

See Supplementary table 1.

### tRNA sequencing of Mouse Brains

Analysis of tRNA abundance, processing and acylation was performed by MesoRNA (Chicago, IL, USA). Paired-end reads were split by barcode sequence using Je demultiplex with options “BPOS = BOTH BM = READ_1 LEN = 4:6 FORCE = true C = false”. Then, fastq files for individual samples were directly used to map against a reference genome of mouse mature nuclear encoded tRNAs, with Arg-TCA-1-1 added, and intron-containing variants of Arg-TCA-1-1 and native Arg-TCT-1-1. Using read 2 only, bowtie2 with “local” mode use used of alignment. From here mapped reads were filtered using a custom script to include only reads where one end is 5’ of position 32 and the other end is 3’ of 37, thus ensuring coverage across the Arg-TCT / TCA anticodon and eliminating ambiguous reads. Abundance figures were derived from counting the number of filtered reads mapping to each reference gene. For modification data, SAM files were processed to WIG files using IGV-2.19.6 command “count” with options “-z 5 -w 1 -e 250 –bases”. WIG files were reformatted with custom scripts from the original MSR-seq pipeline available on github (“https://github.com/ckatanski/MSR-seq“). See Supplementary Table 3.

## RESULTS

### Assessing PTC burden in brain and agnostic rescue in SYNGAP1 and CDKL5

Premature termination codons are a major driver of neurodevelopmental and neurological disorders, **Fig 1A, B**. An analysis from the clinical variant database ClinVar shows that nonsense variants account for roughly 10%-45% of all patients in the 11 genes that were assessed. The highest burdens were observed in *CHD2* (42%) and *SYNGAP1* (35%), followed by *CACNA1A, FOXG1* and *PCDH19* (23%). Across all genes, Arg-to-TGA PTCs were the most common variants, reaching the highest abundances in *SYNGAP1* (54%); *SCN2A* (51%); *SCN1A* (43%) and *CHD2* (42%). A second prominent gene-dependent enrichment involved Gln-to-TAG variants. This class was especially frequent in *SLC6A1* (46%) and *SLC2A1* (41%) and was also present in *CDKL5* (34%) and *FOXG1* (27%), indicating that in these genes a large fraction of nonsense variants arises from Gln-to-TAG substitutions.

Overall, the distribution of nonsense variants is dominated by a small number of recurrent sequence contexts, particularly Arg-to-TGA, with additional gene-specific enrichment for Gln-to-TAG and a limited set of secondary contexts. The submission summary by PTC type showed a similarly skewed pattern, **Fig 1B**. Arg-to-TGA accounted for 34% of submissions and Gln-to-TAG accounted for 17%. Each remaining context contributed 8% or less, forming a long tail of lower-frequency events. Because Arginine CGA-to-TGA events were the most frequent class of PTCs, we next examined local nucleotide composition surrounding patient-derived CGA-to-TGA sites. Outside of the stop codon itself, the flanking sequences showed modest but consistent positional biases rather than a single dominant consensus, **Fig. 1D**. At the position immediately upstream of the CGA codon, Cytosine was enriched (52.6%), with Adenine, Thymine and Guanine each at 15.8%, indicating a preference for Cytosine directly 5’ of the CGA among these sites. At the position immediately downstream, the +1 base was purine-rich (Guanine, 47.4%; Adenine, 42.1%), with Cytosine at 10.5% and no Thymine observed. Across the wider window, base frequencies varied in 5.3% increments (1 of 19 sites), and most positions showed mixed composition without strong fixation. Thus, within this set of patient-derived CGA-TGA variants, enrichment was largely confined to the immediate flanking bases.

### Sup-tRNA rescues translation across patient-derived TGA sites with variable efficiency

To test whether the Arg^UGA^ sup-tRNA supports unbiased suppression across patient-derived TGA sites, we generated cDNA constructs corresponding to each Arg-TGA mutation in *SYNGAP1* and *CDKL5* and fused them to a P2A-NanoLuciferase reporter, **Fig. 1C**.

Constructs were transfected into HEK293T cells either without sup-tRNA (-sup-tRNA) or with a sup-tRNA-Arg-TCA-1-1 expression plasmid (+sup-tRNA), and luminescence was quantified by plate reader 24 hours post transfection.

While all *SYNGAP1-*TGA mutants showed markedly reduced signal without suppressor tRNA (0.7-3.7% of wild type), co-expression of sup-tRNA-Arg-TCA-1-1 increased luminescence for every mutant (37-104% of wild type). This corresponds to 28-82-fold increases relative to the -sup-tRNA condition. Additionally, rescue varied modestly across sites, with the lowest mean signal at R105TGA (1.21×10^6^ a.u.) and near-wild-type recovery for R76TGA and R526TGA (3.25-3.39 ×10^6^ a.u.). Consistent with previous results, *CDKL5-*TGA mutants were reduced without sup-tRNA (0.6-14% of wild type), while co-transfection of the sup-tRNA increased luminescence across all sites. Rescue was weakest at R134TGA (8.2% of wild type), whereas multiple sites (e.g., R59TGA, R550TGA, R559TGA, R970TGA and R981TGA) recovered to approximately wild-type levels. Together, these data show broad suppression across patient-derived TGA sites, with measurable variability between individual mutations.

### Assessing sup-tRNA performance in vitro

Given the potential for an Arg^UGA^ sup-tRNA to repair PTCs in HEK cells, we aimed to develop payload that would be successful for delivery to mammalian brain. Recent effects have made modest improvement in payload designs (22,27) however, in vivo tRNA potency remains a challenge. We therefore tested multiple encoding elements for their ability to produce higher viral vector titers as well as potent payloads with in vivo activity. The variable that we focused on were traditional single-stranded AAV (AAV) and self-complementary AAV (scAAV), as the latter has a smaller double stranded payloads that we hypothesized to be more tolerant of tRNA. We also examined the standard U6 RNA promotor compared to the short native genomic context surrounding the tRNA within its genomic context. Here, we reasoned that the genomic, and potentially the short tRNA intronic, context may provide more efficient expression and tRNA processing. To evaluate the capacity of Arg^UGA^ sup-tRNA constructs to rescue PTCs, we generated vectors encoding the sup-tRNA with or without an intronic sequence, driven by either its native genomic context or an upstream U6 promoter. Constructs were packaged into AAV or scAAV and delivered to HT1080 cells transiently expressing a Nano-Luciferase reporter containing a premature TGA stop codon (NLuc-TGA), **Fig. 2A**. We then quantified reporter rescue using a luminescence-based plate reader assay, **Fig. 2B**.

**Figure 2.**
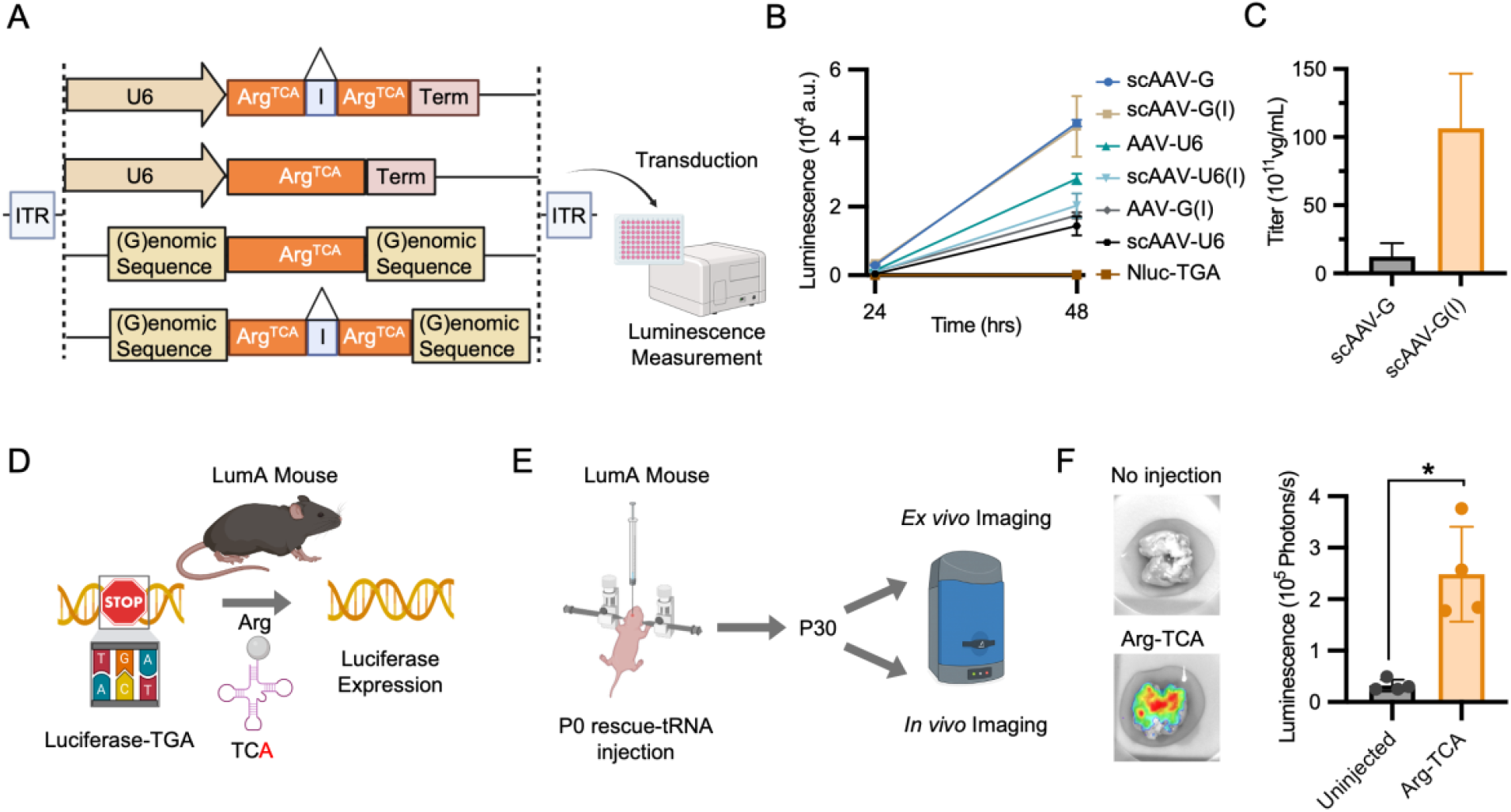
Payload optimization and in vivo assessment. **(A)** Schematic of suppressor-tRNA expression cassettes packaged into ssAAV and scAAV vectors. Constructs are U6-driven or genomically flanked, each including or excluding an intron sequence within the tRNA-Arg-TCT1-1. Vectors were used to transduce NanoLuc-TGA reporter HT1080 cells. **(B)** Luminescence of transduced cells measured at 24 h and 48 h post-transduction demonstrating in vitro PTC suppression efficiency of each construct. **(C)** Viral titers of scAAV-G and scAAV-G(I) measured by qPCR. **(D-E)** In vivo delivery of sup-tRNA constructs into neonatal LumA mice via intracerebroventricular injection. **(F)** Ex vivo luminescence measurement of injected versus uninjected LumA brains confirming localized rescue.

At 24 h post-transduction, all sup-tRNA configurations produced measurable increases in luminescence signal relative to NLuc-TGA controls, although with marked differences in magnitude. Specifically, constructs within a 100 bp genomic context, both with the tRNA intron (scAAV-G(I)) and without the intron (scAAV-G) produced the highest signals, followed by AAV-U6 and AAV-G(I), **Fig 2B**. By 48 h, luminescence signals increased substantially across all conditions, indicating continued accumulation of functional reporter protein over time. scAAV-G and scAAV-G(I) remained the most potent configurations while intermediate signals were observed for AAV-U6, scAAV-U6(I), AAV-G(I) and scAAV-U6; the NanoLuc-TGA reference remained low. These results demonstrate that the Arg^UGA^ sup-tRNA within a genomic context using self-complementary AAV outperforms other configurations in vitro. Interestingly, across three independent production sources (University of Iowa Viral Vector Core; Charles River; PackGene), scAAV intron-containing vectors were consistently higher in titers than those purely within genomic context, **Fig. 2C**.

### Self-complementary AAV enables superior delivery of sup-tRNA in mammalian brain

To assess whether Arg^UGA^ sup-tRNA-mediated rescue could be detected in the brain, we utilized a reporter mouse line that carries a luciferase gene containing an R387TGA premature stop codon at the *Rosa26* locus (28). In this model, luciferase activity provides a direct readout of TGA stop-codon suppression, **Fig 2D**. LumA reporter mice were first injected intracerebroventricularly (ICV) at postnatal day 0 (P0), and brains were harvested at P30 for ex vivo bioluminescence imaging using IVIS, **Fig 2E**. scAAV-G(I) brains exhibited a 7.7-fold increase in signal (*p* < 0.05), indicating sustained reporter rescue in the brain, **Fig 2F**.

After confirming activity of the Arg^UGA^ sup-RNA in mouse brains, neonatal (P0-P1) LumA mice received ICV injections of AAV9 or scAAV9 vectors within the different configurations made, **Fig 3**. At P30, in vivo bioluminescence was quantified by IVIS in live anesthetized mice as region-of-interest (ROI) luminescence and normalized to background luminescence in the LumA mouse line. We used a scAAV-mO2-TGA construct as a negative control for this set of experiments. This approach unlocks a direct comparison of in vivo rescue for payload assessment in mammalian brain. In combination with live animal IVIS transcranial imaging the rescue of the LumA luciferase element can be non-invasively measured and tracked over the lifetime of the animal. At a dose of 7×10^8^ vg per mouse, the mO2-TGA negative control vector showed no increase above baseline luminescence, **Fig. 3A, B**. In contrast, all six tRNA-expressing vectors produced significant reporter rescue when compared to the negative control. The strongest signals were observed for scAAV-G (*p* = 0.0001; *n* = 13) and scAAV-U6 (*p* = 0.0026; *n* = 9) each producing ~3.3-fold increase above background. Constructs containing an intron exhibited more moderate effects, with scAAV-G(I) and scAAV-U6(I) displaying a 2-fold increase in luminescence (scAAV-G(I): *p* = 0.0009, *n* = 9; scAAV-U6(I): *p* = 0.0485; *n* = 9), while single-stranded AAV vectors yielded intermediate rescue of ~2.8-folds over background (AAV-U6: *p* = 0.0074, *n* = 9; AAV-G(I): *p* = 0.0038; *n* = 10).

**Figure 3.**
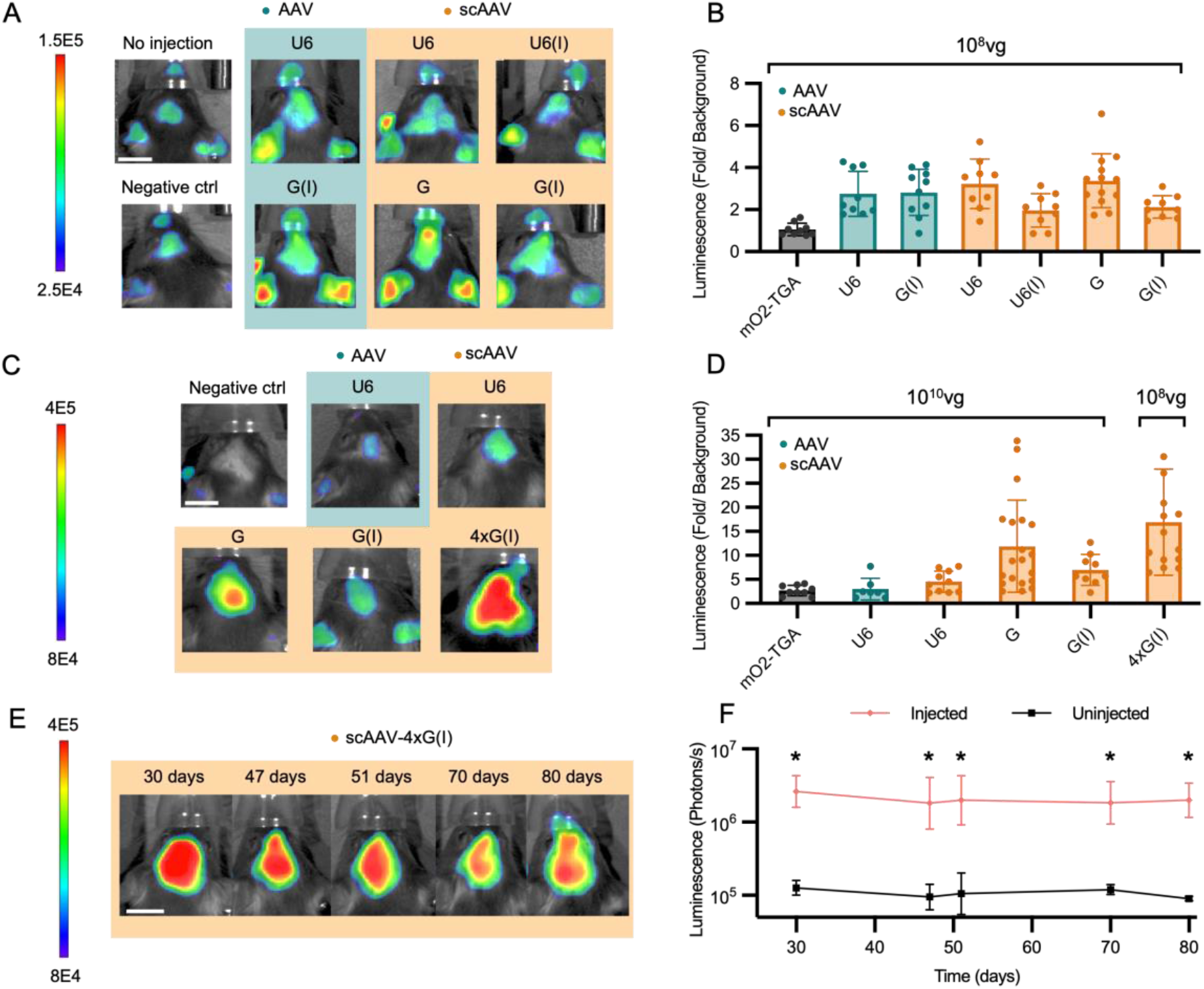
Potent and durable UGA rescue in brain with self-complementary AAV and minimal genomic context. **(A)** Representative transcranial IVIS images and quantification of LumA mice injected with a low dose of suppressor tRNA constructs (7×10^8^vg) **(B**) Luminescence rescue (fold/background) for each injected condition at 7×10^8^vg. Scale bar: 1cm. **(C-D**) Genomic context provides highest rescue and multiplexing to 4X tRNA copies enables similar rescue at 14-fold lower viral dose of 1×10^10^vg. **(E-F)** Longitudinal IVIS imaging demonstrating durable nonsense suppression following scAAV-4xG(I) delivery over 30-80 days post-injection. Data are shown as mean ± SD with individual biological replicates.

Increasing the dose to 1×10^10^vg further enhanced reporter rescue, **Fig. 3C, D**. While the scAAV-mO2-TGA negative control showed a modest signal in luminescence (*n* = 9), self-complementary tRNA-expressing vectors with the genomic sequences produced substantially higher luminescence levels. Relative to this control, scAAV-G produced the largest increase (12-fold over background; *p* = 0.002; *n* = 20), followed by scAAV-G(I) (7-fold; *p* = 0.016; *n* = 9). Meanwhile, scAAV-U6 (*p* = 0.14; *n* = 10) and AAV-U6 (*p* = 0.99; *n* = 7) showed no significant differences than the negative control at the tested dose, yielding 4.5-fold and 2.9-fold increases over background, respectively. Notably, at the lower dose of 7×10^8^ vg per mouse, increasing the copy number of the tRNA to four, while keeping the genomic and intron sequences, outperformed the higher dose of 1×10^10^ vg per mouse. scAAV-4xG(I) yielded over 16-fold over background (*p* = 0.001; *n* = 14), exceeding the mean signal of all high-dose constructs in this dataset, **Fig. 3C, D**. Thus, the small size of the tRNA in the payload opens up the possibility of tRNA multiplexing and lower viral doses.

### scAAV9 tRNA delivery provides durable rescue

To determine whether tRNA-mediated rescue persists over time in vivo, we next assessed the durability of the most effective tRNA construct, scAAV-4xG(I), using longitudinal bioluminescence imaging in LumA mice, **Fig. 3 E, F**. Following ICV neonatal injections, reporter signal was quantified between postnatal day 30 (P30) and P80 and compared to age-matched uninjected controls. Luminescence in injected mice remained consistently elevated, showing a 17.25-24.17-fold difference between groups across time points (*p* < 0.0001).

### In vivo tRNA-Arg-TCA-1-1 isodecoder abundances, charging profiles, and processing

There are 28 human arginine tRNA genes and it is possible that viral expression of a sup-tRNA may alter the expression balance and/or tRNA acylation efficiency. Indeed, all Arg tRNA are serviced by a single AARS. Thus, the unintended effect of Arg^UGA^ sup-tRNA expression is the competition for AARS activity. To determine whether neonatal scAAV-G(I) delivery alters the endogenous tRNA landscape in brain, and to confirm expression and aminoacylation of the Arg-TCA-1-1 sup-tRNA, ICV injected LumA mouse brains were harvested at P30 and tRNA were extracted for full sequencing and analysis. By analyzing the total mitochondrial (mt) and cytosolic (Cyt) tRNA read counts, sequencing data revealed that both, injected and uninjected brains, expressed similar abundances: cytosolic tRNAs comprised 95.6% and 94.1% of the total tRNA respectively, while mitochondrial tRNAs accounted for 4.4% and 5.9%, **Fig. 4A**.

**Figure 4.**
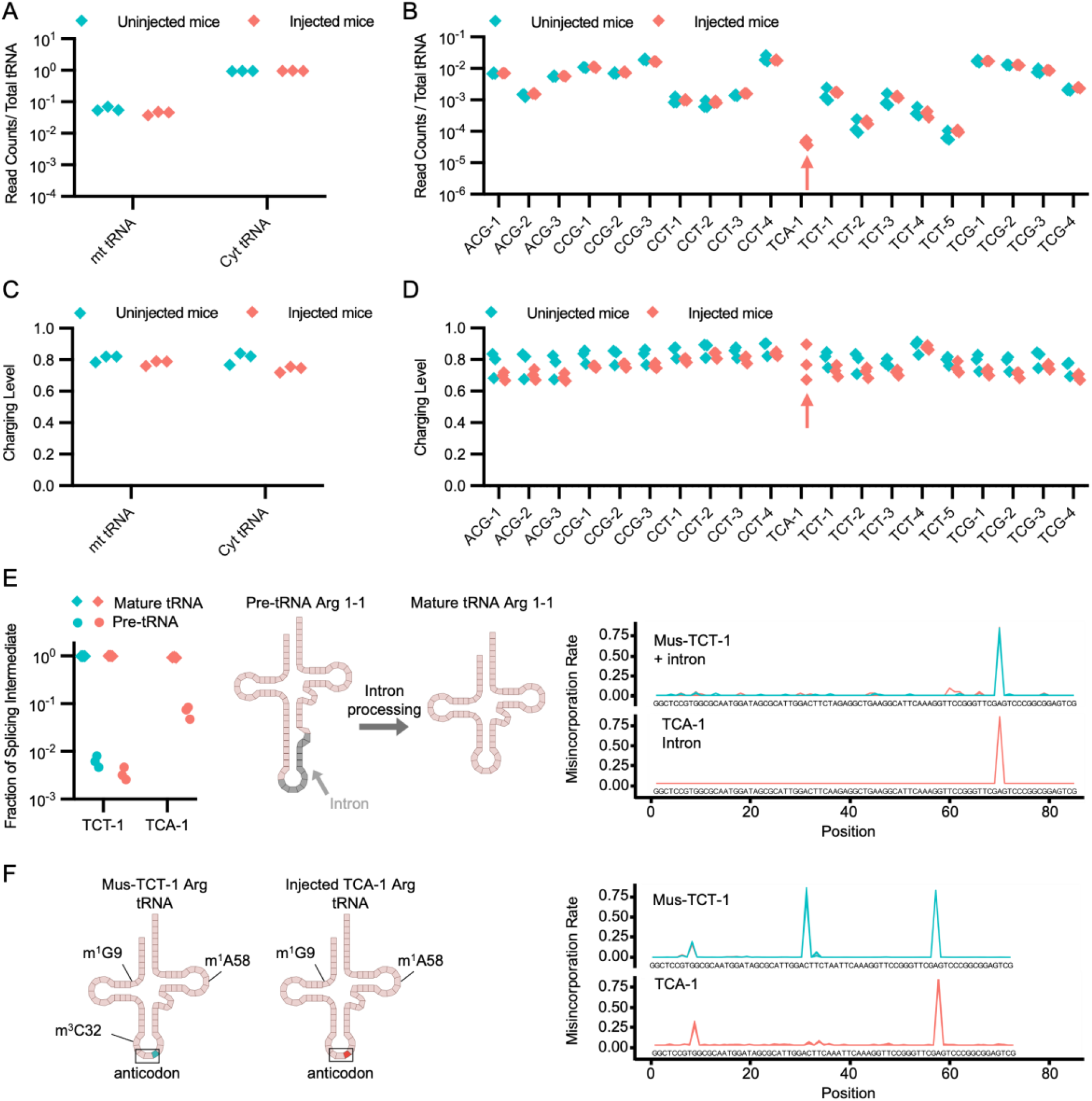
Brain tRNA sequencing shows preserved endogenous tRNA abundance, aminoacylation, and processing following sup-tRNA delivery. **(A)** Global tRNA abundance in uninjected and sup-tRNA-injected mouse brains, separated into mitochondrial (mt) and cytosolic (Cyt) tRNAs. **(B)** tRNA-Arg isodecoder-level abundance in both conditions showing detection of tRNA-Arg-TCA-1 only in injected brains with no apparent changes across endogenous Arg-tRNAs. **(C)** Aminoacylation levels of mt and Cyt tRNA. **(D)** Charging levels of tRNA-Arg isodecoders confirm charging of tRNA-Arg-TCA1-1 in injected brains. **(E)** Left: Fraction of mature versus premature parental and sup-tRNAs in uninjected and injected brains. Right: Misincorporation profiles indicate similar intron processing in both **(F)** Arg-tRNAs. Misincorporation profiles comparing parental tRNA-Arg-TCT-1-1 to tRNA-Arg-TCA-1-1 in uninjected and injected brains, arrows denote the modification signature sites.

Within the cytosolic Arg tRNA isodecoder dataset, all species showed similar fractional abundances between the two groups, with the sup-tRNA readily detected in the injected mouse brains and comprising only 1.2% of the tRNA-Arg-TCT anticodon family, **Fig. 4B**.

Additionally, the charging fractions for the global tRNAs as well as the Arg tRNAs in both groups were not different, with Arg-tRNA-TCA-1 ranking right above the median, **Fig. 4C, D**. To confirm that the sup-tRNA undergoes canonical maturation in vivo, we quantified the relative abundance of mature versus intron-containing (premature) tRNA species in injected mouse brains. For tRNA-Arg-TCT-1-1, the mature fraction was near-complete in both uninjected and injected brains, consistent with robust endogenous processing, **Fig. 4E**. Similarly, tRNA-Arg-TCA-1-1 was efficiently processed in the injected mice, as indicated by the predominance of the mature tRNA species following intracranial delivery.

We next compared misincorporation-rate profiles from tRNA-Arg-TCT-1-1 and tRNA-Arg-TCA-1-1 across conditions, **Fig. 4F**. In both injected and uninjected brains, peaks were observed at positions corresponding to m^1^G9 and m^1^A58 for both Arg tRNA species. This indicates that the delivered tRNA is recovered by tRNA-seq and undergoes canonical maturation and modification steps in vivo. In contrast, a signature at position 32 (consistent with m^3^C32) was present for tRNA-Arg-TCT-1-1 but absent for tRNA-Arg-TCA-1-1. Therefore, the Arg^UGA^ sup-tRNA is processed as a bona fide tRNA substrate in brain while retaining an anticodon-dependent modification state consistent with its engineered identity.

## DISCUSSION

Here we describe a programmable method using human suppressor tRNA which enables a general strategy to repair UGA PTCs in mammalian brain. A primary concern of any read-through strategy is the potential interactions with native stop codons which terminate translation. For instance, small molecules read-through agents (e.g. Ataluren) promote mis-encoding at the site of the PTC as well as elsewhere in the proteome (12), yet are deemed safe for clinical use (29). By contrast, sup-tRNA encode the correct amino acid at the PTC – a key feature for proteins with low tolerance for missense variants – and appear to have minimal interactions with native stops (18,24,30). Possible mechanisms for this observation have been previously summarized (17) and suggest that engineered tRNA display a “stop codon” bias for PTC variants over native 3’ stops due to the genetic context of native stop codons (31,32). That is, eukaryotic mRNA circularization orchestrates a ribonucleoprotein (mRNP) complex between the 3’ poly-binding protein (PABP), the 5’ RNA-binding protein elF4E, and a ternary complex of release factors (eRF1, eRF3, GTP). There are likely many unresolved details in this mechanism and additional work is required to understand how this ribo-protein complex works in concert to minimize sup-tRNA read-through. An additional challenge for the development of a tRNA-based therapy is the possibility that expression of a sup-tRNA could alter the processing or expression of other tRNAs from the same family (e.g. Arginine). The data generated here from virally delivered tRNA in brain suggest that, at the doses delivered, tRNA acylation, methylation and expression are unchanged. However, future studies will be needed to better understand the tRNA expression threshold of these observations. Taken together, these data show that there is a therapeutic window of sup-tRNA expression that promotes PTC repair in vivo, yet does not affect native stop codons or tRNA processing.

With this apparent safety profile in mind, we examined tRNA-bearing payload designs in the brains of P0 mice using intracerebroventricular injection of viral vectors. The reason for this experimental approach was two-fold. First, many of the neurodevelopmental disorders shown in **Figure 1** would benefit from early intervention, as it is unknown if there are therapeutic windows where a given repair strategy will be most effective. It is reasonable to believe that earlier treatments are, generally, better (33,34). Second, the neurodevelopmental changes in the mammalian brain between birth and adolescence are marked by extreme molecular, cellular, and structural growth, making this an especially fragile and fraught period for gene regulation. By delivering our most potent tRNA payload, a 4X Arg^UGA^ sup-tRNA based on the natural human tRNA with highest activity, this approach serves as a tolerance test of the method as the tRNA would be present during the many crucial milestones of brain development. Notably, we observed no obvious changes to mouse behavior or physiology, although additional studies will be needed to expand this analysis to subregions of the brain and to perform more detailed behavioral assessments.

In conclusion, this work provides a safety threshold for sup-tRNA in vivo, a means for efficient viral production and assessment of biological activity with the LumA mouse model, and methods to quantify of target interactions. Given the unmet need from the many currently untreated genetic diseases caused by PTC variants, this study brings the possibility of a first-in-class therapeutic tRNA strategy one step closer to the clinic.

## ACKNOWLEDGEMENTS

We thank our lab members and colleagues for their helpful discussions and feedback on the manuscript.

## AUTHOR CONTRIBUTIONS

Conceptualization, A.A.S., L.G., C.A.A; Methodology, A.A.S., L.G., C.A.A; Investigation A.A.S., L.G.; Data Analysis and Visualization, A.A.S., L.G; Writing – Original Draft; A.A.S., C.A.A Writing – Review and Editing; A.A.S., L.G., C.A.A

## SUPPLEMENTARY DATA

Three supplemental tables are provided for cloning details, sequences and tRNA sequencing files.

## Supplementary Data statement

Supplementary Data are available at NAR online.

## DATA AVAILABILITY

All relevant data are included within the article and its Supplementary Information files. Requests for further information and resources should be directed to and will be fulfilled by the lead contact, Christopher A Ahern.

